# A model of marmoset monkey vocal turn-taking

**DOI:** 10.1101/2023.10.07.561358

**Authors:** Dori M. Grijseels, Daniella A. Fairbank, Cory T. Miller

**Author notes:** Direct Correspondence to or.

## Abstract

Vocal turn-taking has been described in a diversity of species. Yet a model that captures the various processes underlying this social behavior across species has not been developed. To this end, here we recorded a large and diverse dataset of marmoset monkey vocal behavior in social contexts comprising one, two and three callers and developed a model to determine the keystone factors that affect the dynamics of these natural communicative interactions. While a coupled oscillator model failed to account for turn-taking in marmosets, our model alternatively revealed four key factors that encapsulate much of patterns evident in the behavior, ranging from internal processes, such as the state of the individual, to social context driven suppression of calling. In addition, we show that the same key factors apply to the meerkat, a carnivorous species, in a multicaller setting. These findings indicate that vocal turn-taking is affected by a broader suite of mechanisms than previously considered and that our model provides a predictive framework with which to further explicate this natural behavior in marmosets and for direct comparisons with the analogous behavior in other species.

## Introduction

Reciprocal vocal exchanges, such as chorusing, duets and conversations, are common across a diversity of vertebrate species, including humans,^1^ monkeys,^2^ mice,^3,4^ and birds.^5^ These interactions are governed by a system of social rules, some of which are species-specific and others shared across taxa.^6,7^ A classic example of the latter is turn-taking; the behavior of speakers alternating the timing of their successive calls in a sequence. The widespread occurrence of turn-taking offers the opportunity to leverage similarities and differences in this behavior to elucidate the various underlying mechanisms of this dynamic behavior.^6^ Whereas core sensory and motor mechanisms are inherent to turn-taking, as the conversationalists must perceive and produce the social signals, speakers must also attend closely to all conspecifics in the scene and adapt their behavior in response to ongoing contextual changes in the social and ecological landscapes. Turn-taking, afterall, emphasizes the fact that conversations reflect a social interaction that is not only supported by sensory and motor processes but a suite of complementary cognitive processes that are each necessary for these coordinated exchanges to manifest. Explicating how each of these processes complement each other is crucial to understanding potential mechanistic differences that support seemingly similar behaviors across species.

The common marmoset (Callithrix jacchus), a small New-World primate, engages in vocal turn-taking during antiphonal conversations, a vocal interaction involving the recriprocal exchange of the species-typical phee calls that emerges when animals are visually occluded from each other.^8–14^ As with turn-taking in other species, marmoset antiphonal conversations are characterized by a lack of overlapping phee calls^8,12,15^ and coordinated call timing and duration.^11,12,16–18^ Recently it was proposed that, like humans,^19^ marmosets act as coupled oscillators such that the timing of phee call production between two marmosets will become coupled and entrained during extended conversations.^8^ However, several features of marmoset vocal behavior are not accounted for by this hypothesis. Marmosets, for example, will alter their call timing to an individual call, depending on its features,^20^ and do not reliably respond to the same call under similar circumstances.^18^ There is also evidence that internal factors can affect marmoset vocal behavior in some contexts^21^. These findings suggest that turn-taking is affected by a number of different processes and that a more comprehensive model of marmoset conversations is needed that fully encapsulates the various factors affecting turn-taking in this primate species.

Here, we sought to characterize turn-taking during marmoset conversations by first recording a large and diverse representative dataset (42 subjects; 11,614 phee calls) and then to use these data to develop a novel model of the natural dynamics of turn-taking. We first tested and failed to replicate the coupled oscillator hypothesis for the larger population of marmosets tested here, instead finding that respective call timing between marmosets in these exchanges was stochastic. Following a thorough analysis of marmoset vocal behavior, we propose a novel model, based on internal and external states, that describes conversational dynamics across various paradigms from single, pairs and trios of marmosets. Overall we show that while a simplified model based on volubility factors heavily in characterizing their behavior in some contexts, social cognitive processes modulate vocal behavior, particularly in larger groups reflective of the communication networks typical of marmoset societies.

## Results

### Marmoset monkeys do not act as coupled oscillators during turn-taking

To examine the dynamics of marmoset conversations, we employed a two monkey paradigm consistent with previous studies^12,22^ (S Fig 1A). Two marmosets were placed into test boxes approximately 4m apart on opposite sides of a soundproofed room with an opaque cloth occluder positioned at the midpoint to eliminate visual access between the animals. Directional microphones placed in front of each animal recorded the vocalizations of each monkey, which we extracted for analysis.

**Figure 1:**
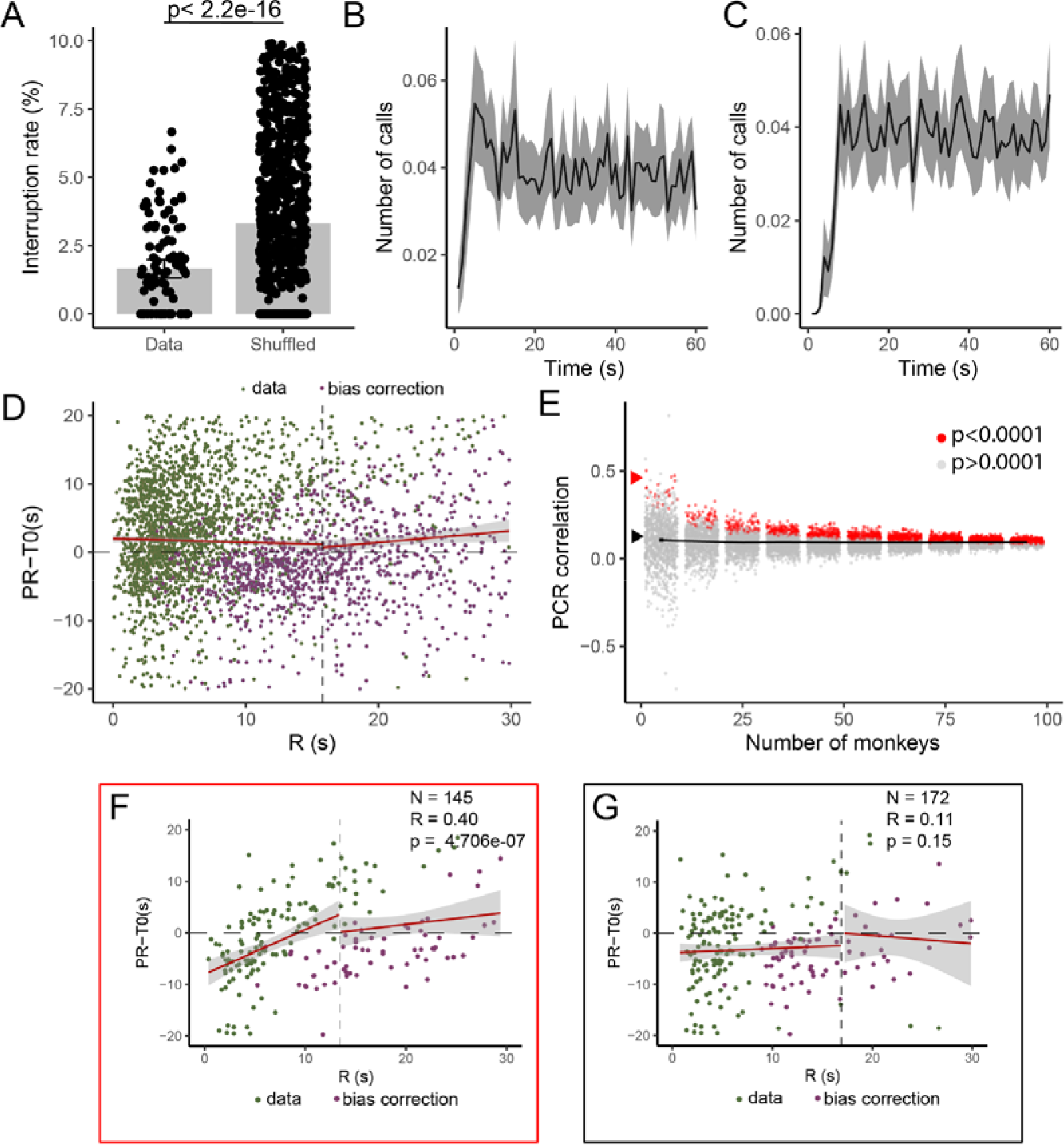
Replication of analyses showing monkeys do not act as oscillators. (A) Percentage of calls interrupted compared to a shuffled control. Proximate measures (replicated from Takahashi et al.) of crosscorrelation (B) and autocorrelation (C) across all sessions. (D) PRC plot of the vocal exchanges. (E) PRC correlation values at different subset sizes, red dots indicate datasets with a significant correlation (p<0.0001). Red arrow points to dataset showing in (F), black arrow shows (G). (F) PRC of a single 10 monkey subset (N=145 calls), showing a possive PRC correlation. (G) PRC of a representative median 10 monkey subset (N=172 calls), showing a non-positive PRC correlation.

A primary aim of this study was to record a large corpus of recording sessions from animals across different ages, sex and social relatedness to accurately characterize the species’ conversational dynamics. We recorded the vocal behavior of 42 different marmoset monkeys, including 59 male-female pairs, 21 female-female and 27 male-male pairs (S Fig 1B). 25 pairs (23%) were cagemates, out of those 7 were bonded pairs, 14 were parent-sibling pairs and 4 were siblings (S Fig 1C). On average the age of the monkey at recording was 1179 days (range 260-2770 days), we included pairs with a range of age differences (SFig 1D), and recorded at varying times of the day (SFig 1E). While duets are limited to paired males and females in many species,^23–26^ conversations in marmosets occur between individuals of various ages, ranks and sexes suggesting that differences may emerge in patterns of this vocal behavior along these different social categories. However, contrary to previous studies^12,17^, we found no effects of sex, age, cage mate status, or time of day on the response rate (SFig 1F-I).

The current leading theory for marmoset turn-taking dynamics is that monkeys in a pair act as coupled oscillators.^8^ This hypothesis states that monkeys call in an anti-phasic manner: when one monkey talks the other is quiet and vice versa, and the timing of one animals response determines the timing of the subsequent response in the partner. This oscillatory dynamic underlies the coordination of vocalizations during a conversation. Using the dataset collected here, we applied the identical analyses^8^ to test whether the coupled oscillator theory generalised to a larger population of marmosets. We first confirmed that in agreement with Takahashi et al.^8^, we observed a decrease in interruption rate compared to a shuffle control, indicating that marmosets actively avoid overlapping calls (i.e. engage in turn-taking; Figure 1A). Unlike previous results^8^ some overlapping calls did occur between monkeys in our dataset. While interruptions were rare, only accounting from 234 calls out of 11450 calls, 61.7% of our sessions had at least one overlapping call. We hypothesized this might be due to the inclusion of younger monkeys,^9^ however, there was no significant correlation between age and the number of interruptions (SFig 1A).

We next tested whether our monkeys showed an oscillatory pattern in timing of responses after a partner call by replicating the analysis performed by Takahashi et al.^8^. Unlike their findings, we observed no oscillatory pattern in our data (Figure 1B), nor was an oscillatory pattern evident in a proximate measure forthe autocorrelation of the monkeys (Figure 1C). This indicates that when a larger dataset is tested the marmosets do not act as oscillators, a necessary requirement for a coupled oscillator dynamic. To demonstrate that marmoset behaved as a coupled oscillator in their conversations, Takahashi et al.^8^ showed a positive correlation between the time in between two calls by the same monkey (PR), corrected for the average time between two calls by that monkey when not in conversation (T0), and the time between a monkey and its partner’s call (R). Here we find no such correlation between R and PR-T0 (Figure 1D). In addition, the PR-T0 in this analysis does not cross the 0 line, indicating that a long interval by monkey 1 did not result in a shorter response latency by monkey 2. These findings demonstrate that turn-taking in marmoset conversations does not follow coupled oscillator dynamics.

Since our results differ so drastically from the previous findings, we next tested whether any smaller subsets of our data of a similar size as the previous study^8^ would yield results that matched these findings. We randomly sampled subsets of our data and performed the PRC analysis on these subsets and the PCR. We found that the correlation coefficient averaged around -0.01 regardless of dataset size, but there was large range in correlations especially when only including 5 sessions (- 0.74-0.81, Figure 1E), a similar sample size to the previous study^8^. While this analysis revealed that outlier subsamples (defined as any samples with a p<0.0001 and r>0, total of 25 samples (2.5%)) from our dataset could replicate the previous effects^8^ (Figure 1F), the vast majority of sessions (97.5%) resulted in a non-significant, or significant negative correlation (Figure 1G). Together, these results show that although the results from Takahashi et al.^8^ may reflect a small subpopulation of marmoset vocal behavior, a coupled oscillator mechanism is not representative of marmoset conversations as a general rule. While turn-taking reflects a degree of coordination between individuals, our analyses indicated that the respective call timing of each monkey during these exchanges is stochastic and that considerations of factors other than the response timing of the other conversationalist are needed to more completely understand this natural vocal behavior.

### Turn-taking probability and timing is most strongly predicted by marmoset volubility

To develop a model that more accurately characterized marmoset vocal turn taking, we first sought to thoroughly quantify different facets of the vocal behavior and call acoustics and test if any features were predictive of whether and when a marmoset would call in a conversation. For each call, we calculated 12 parameters to first evaluate what parameters affect whether a monkey will respond or not (Figure 2A). These were the calling rate (average prior to the call) of both the monkey and the partner, the previous call’s loudness, partner intercall interval (partner ICI), start frequency, end frequency, maximum frequency, slope of drop in frequency, number of pulses, length of the call, length of the last pulse, and the number of consecutive calls in the current conversation (see Methods for detailed descriptions).

**Figure 2:**
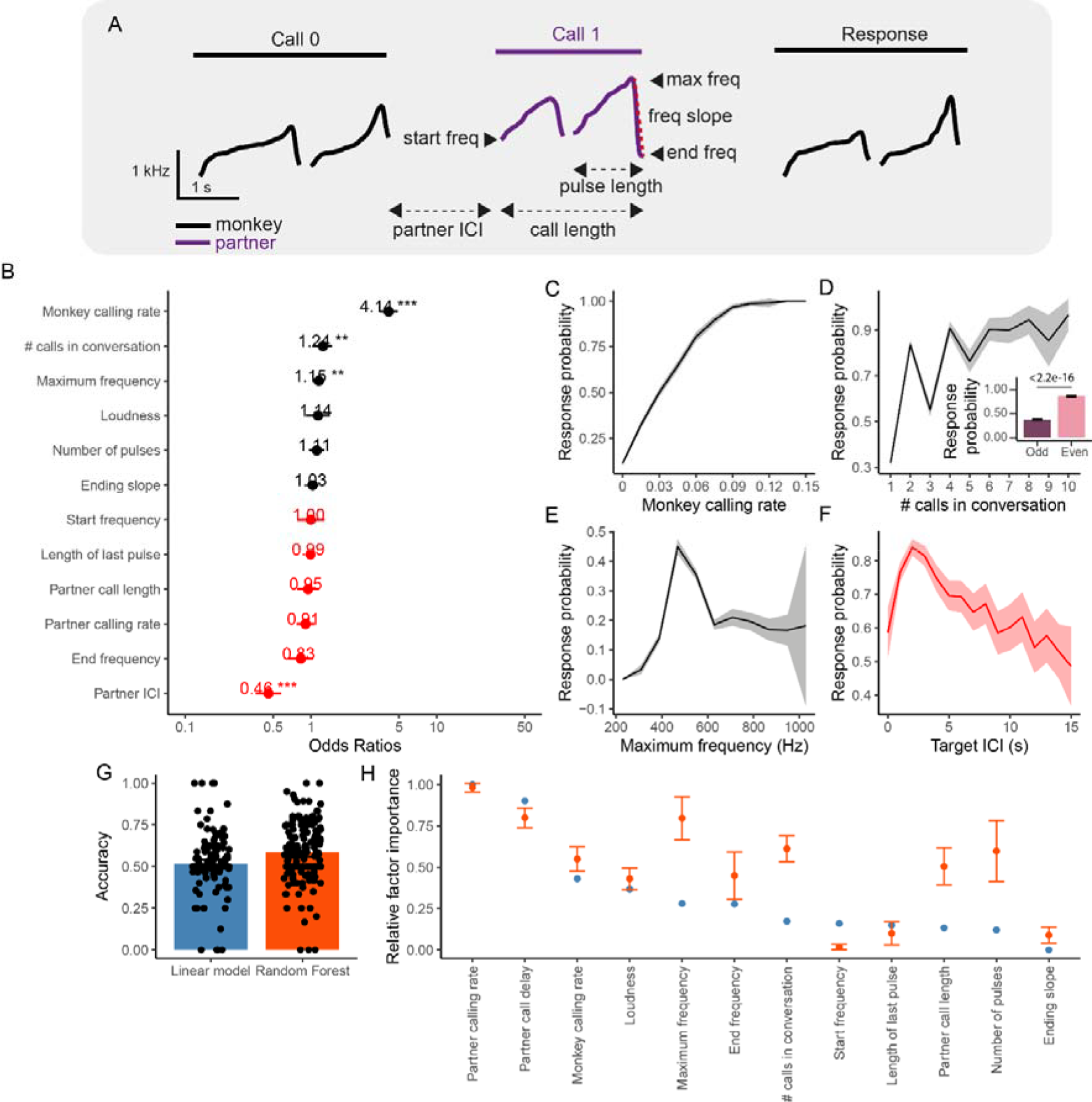
Factors affecting the response probability of marmosets. (A) Example call interaction showing how the various call parameters are measured. (B) Odds ratio of all 12 parameters as resulting from the Generalized Linear Mixed-effects Model. Effect of (C) monkey calling rate, (D) position of call in the conversation, (E) maximum frequency of the call, and (F) the target ICI on the response probability. (G) Accuracy of a linear model and random forest classifier on predicting whether a call will occur. (H) Relative factor importance (normalized) of the linear model (blue) and random forest (orange).

We fit a Generalized Linear Mixed-effects Model (GLME) to the marmoset vocal behavior data with 12 parameters as our fixed-effects and monkey and partner ID as the random effects, and whether or not there was a response as the dependent factor, using the lme4 package in R (marginal R^2^=0.47).^27^ This showed 4 significant factors (Figure 2C): the mean calling rate of the monkey (Figure 2C), the position of the call in the conversation (Figure 2D), the maximum frequency of the previous call (Figure 2E) and the previous ICI (Figure 2F). Interestingly, although there was a general trend of the call probability increasing with the position of the previous call in the conversation, a sawtooth pattern emerged with odd numbered calls in the sequence exhibiting a lower probability than even numbers (Figure 2D), a pattern that resulted in a statistically significant difference (Figure 2D, inset).

Next, we quantified features of the vocal response latency, the main factor predicted by the coupled oscillator. Similar to previous studies,^16^ most marmoset vocal responses recorded in the current study occurred ~3 seconds after the offset of the conspecific call (SFig 2B). We combined all factors into a Linear Mixed-effects model, with the same fixed and random factors as previously described, and response delay as the dependent factor, and found that only the calling rate of the responding monkey significantly affected the response delay (SFig 2A). In other words, a marmoset that called at a higher rate was more likely to respond to a conspecific’s call more quickly (SFig 2C), while the call delay does not affect the response delay, as would be the case in a coupled oscillator system. Overall, this suggests that response latency during turn-taking at the level of single calls cannot be accuretaly predicted with the tested factors and is largely stochastic.

These analyses indicated a relationship between the aforementioned four factors and marmoset response probability. However, this finding does not necessarily mean that these factors can predict whether an individual call within a session will receive a response by the conspecific partner. To test this issue further, we trained two classifiers: a logistic regression and a random forest classifier using the 12 factors described above, as well as an additional factor describing whether the call was an odd or even call in a conversation, on data counterbalanced per session. The logistic regression was not able to predict a response significantly above chance (Figure 2G, p=0.072, student’s t-test), suggesting no linear relation between the tested factors and the response probability, while the forest classifier did predict significantly above chance (Figure 2G, p= 1.434e-07), indicating a non-linear effect is at play.

However, when considering the predictor importance based on the out-of-bag error — i.e. predictor importance estimates by permutation for the random forest classifier — the partner’s calling rate is the top predictor (Figure 2H). Indeed, when we exclude the responding monkey’s calling rates from the classifiers, the random forest classifier only has an accuracy of 58% (SFig 2D). Though still significantly above chance, this finding suggests that the volubility of each marmoset, rather than the specific acoustic properties of individual call or vocal behavior, is the main factor in our model determining whether or not a monkey will respond to a given call. In other words, a marmoset exhibiting high volubility in a session will be more likely to produce a call in response to hearing a conspecific’s call, and a marmoset will receive more responses when conversing with a partner with a high volubility. The large remaining unexplained variance in our models may have either of two causes. Either we did not measure all relevant behavioral characteristics that covary with the probability of a vocal response (i.e. head-turning,^14^ arousal,^28^ attention, etc) in the current study, or a truly stochastic process is taking place to determine whether an animal will respond.

### A novel model of marmoset turn-taking

Here, we propose a model of marmoset vocal turn-taking that combines an individual’s volubility with several internal and external modulators, which together determine a response probability. Our previous analyses showed that both social (SFig 1F-I) and call-based (Figure 2, SFig 2) factors minimally affect a monkey’s calling behavior in dyads, while individual volubility strongly predicts a monkey’s calling and response rates (SFig 1J-L, Figure 2B,C,H, SFig 2). We based the modulators on both the data presented here, as well as existing studies in the relevant literature, including previous studies showing the effect of various internal^28,29^ and external^20,30^ factors affecting marmoset calling, and relevant modelling work.^20,21^

During the recording sessions performed here the majority of marmosets showed fluctuations in their calling rate over time (see example SFig 3A), likely reflecting changes in arousal^28^, behavioral context^20,29^, and other internal processes, which we jointly label ‘behavioral state’. Analyses testing whether these internal state fluctuations covaried between each pair of monkeys found no significant difference compared to a shuffled control (SFig 3B). A linear model was used to simulate novel fluctuations in calling by training it to predict the calling rate in each 30 second time bin based on the previous 5 time bins (Figure 3A). This was then applied over the test session to iteratively produce novel session-wide internal state fluctuations for each monkey, which was then input into the model (see Methods for details). In addition to independent state-wide fluctuations, an overall trend in the monkeys’ calling rate decreasing over time was evident and labelled as ‘volubility decay’. The actual source(s) of this decay for a given session is difficult to determine, however, we believe it may reflect a combination of arousal, interest, motivation, and unknown other factors. Further studies could potentially separate out these various factors, leading to a more detailed model of this effect. Given the current data, however, the most accurate representation was to fit an exponential decay function (Figure 3B dashed line) to model the decreasing volubility of the monkey over the span of a session.

**Figure 3:**
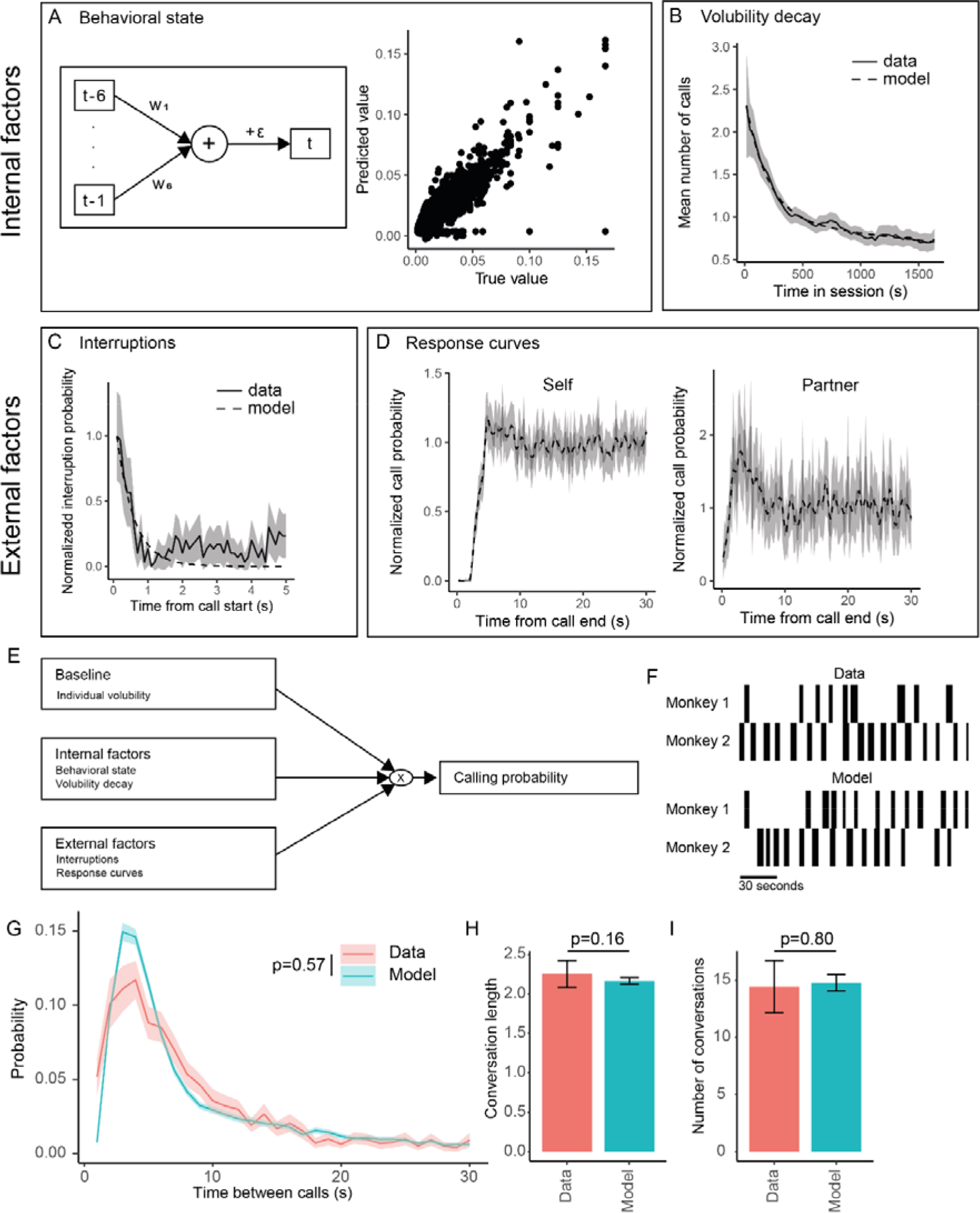
Overview of novel marmoset phee communication model. First row represent the internal factors (A) internal state, as modelled by a linear model with the structure illustrated left, which can predict the state value based on the preceding 5 timepoint (right). (B) Arousal, as modelled by an exponential decay across the session (dashed). External factor include (C) interruption curve, as modelled by an exponential decay (dashed), and reponse curves, obtained from the data. (E) shows the full model schema, with the factors resulting in a calling probability for each timepoint. (F) The first 3 minutes of conversations from a real datasets (top) and the corresponding modelled dataset (bottom). (G) the intercall interval (ICI) of the data (pink) compared to the model (blue). The mean conversation length (H) and number of conversation (I) of the data (pink) compared to the model (blue).

Turn-taking is affected by at least two key external factors related to the vocal behavior of the other monkey in the conversation. First, it is well established that marmosets largely avoid interrupting conspecific calls^8,9,12^ (Figure 1A). Analysis of the timing interruptions in the current dataset found that the majority of calls were interrupted within the first 0.5s (Figure 3C), likely representing covocalisations, where both marmosets initiate a call at approximately the same time, with a rapid decline over time. This was modelled using an exponential decay, representing the rapid decline in likelihood of calling by a monkey while the partner is vocalising. Second, the periodicity of turn-taking in marmoset converations with phee call is notably slow relative to other species.^19,31–33^ We analyzed the average call probability of the marmoset producing a call immediately following the offset of its previous call and revealed a refractory period of about 2.2s (Figure 3D). While a small portion of this duration is the result of physical constraints, i.e. the monkey needs to take in a breathe before being able to produce a second phee call,^28^ much of this period is likely active suppression of vocal production to allow for the other conspecific turn to respond. Following this refractory period, a rapid rise in call probability occurred before returning back to baseline around 4.6s after the offset of the previous call. The partner marmoset exhibits a complementary pattern of vocal behavior. This individual exhibits an immediate increase in calling probability after hearing a call, peaking at 3.0s before returning to baseline around 7.6s after the end of the call, consistent with previous findings.^16^

Our novel turn-taking model combines all of the four factors described above - behavioral state, volubility decay, interruptions, and response curves - to represent the calling probability over time in the following manner. First, each monkey’s baseline volubility and the arousal and internal state factors were used to first calcalute the session-wide probabilities. Next, we simulated each time step, determining whether a phee call is produced based on the calling probability at that time point for each marmoset, and applying the appropriate external factors when a call is emitted. Using this procedure, we were able to simulate marmoset conversations and generated ‘turn-taking’ using a 5-fold cross-validation paradigm. For each fold, 20% of the dataset was used as a test set, basing the response curves after a call only on the remaining 80%. To validate the model from the test set, the calling rate of our actual sessions was input into the model and the resulting dataset analyzed (see Figure 3F for example interactions from the data and model).

Analyses showed that the resulting ICI probability density distribution, a measure used previously to determine model accuracy ^8^, was not significantly different from the data ICI (Figure 3G). Moreover, no significant differences were evident between the model and the data for the number and length of the conversations emerging from the models (Figure 3H,I). To test the individual effects of the four factors - behavioral state, volubility decay, interruptions, and response curves – each was excluded separately and the changes to the conversation dynamics quantified. Analyses revealed that that only the response factor significantly affected the ICI distribution (SFig 4A) and the length (SFig 4B) and number (SFig 4C) of the conversations. Lastly, the same GLME was applied to the modelled data, excluding the call length and frequency factors as these were not varied in the model, which showed significant effects of monkey call rate, partner ICI and conversation number (SFig 4D-H). In addition, the sawtooth pattern that was observed in the effect of the position of the call in a conversation on the response probability was evident in the modelled data (SFig 4F). Neither the partner ICI, nor the conversation number effects were explicitly modelled, suggesting these results in the data were emergent from the conversation structure and calling rate, rather than causative results.

**Figure 4:**
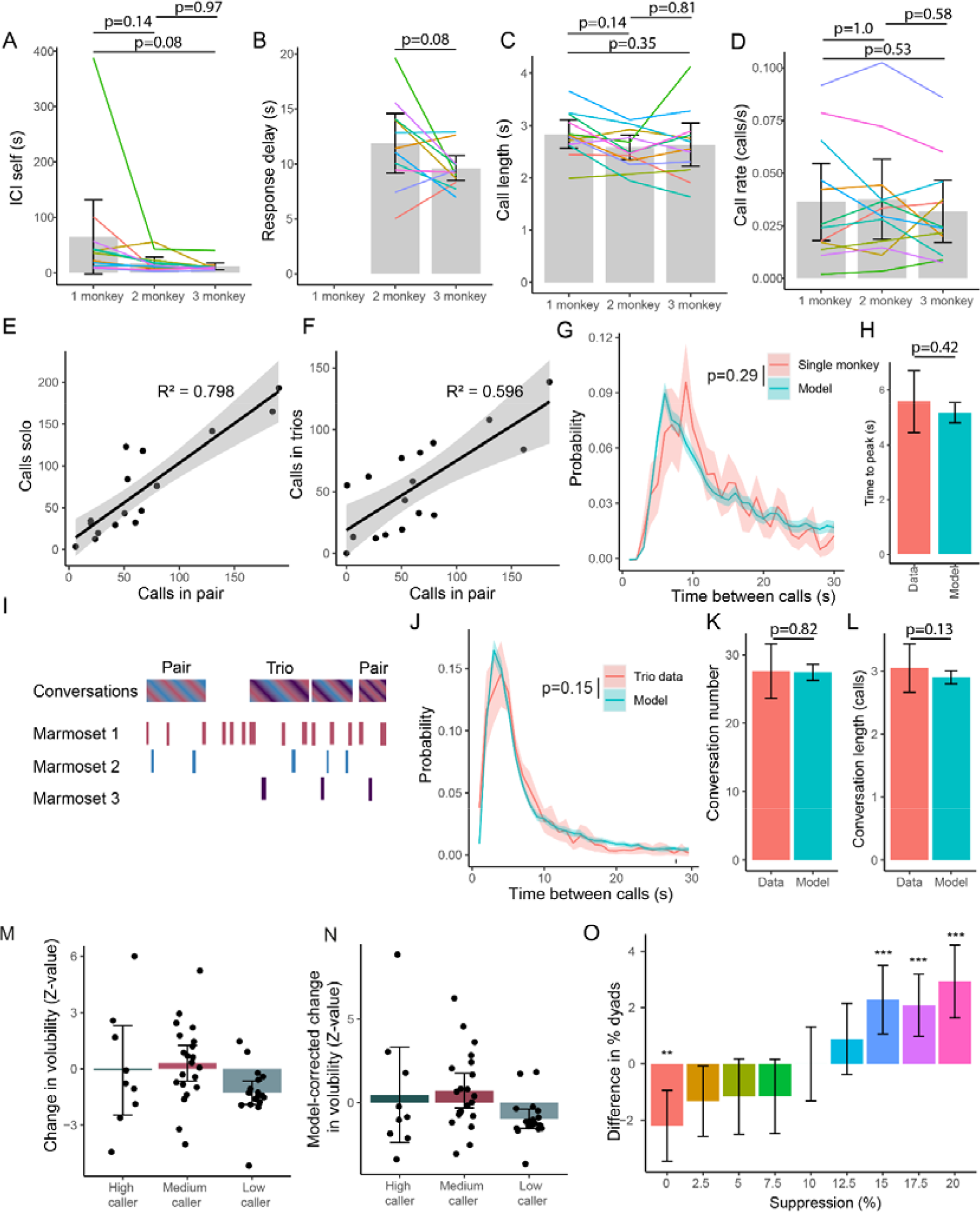
Novel marmoset communication model applied to novel paradigms. (A) ICI between consecutive calls by the same monkey across the three paradigms. (B) Response delay between calls of monkeys engaging in conversation. (C) Mean call length of monkey across the paradigms (D) Calling rate of monkeys across three paradigms. (E) The mean calling rate of individual monkeys in the paired condition compared to the solo condition. (F) The mean calling rate of individual monkeys in the paired condition compared to the three monkey condition. (G) The intercall interval in the data (pink) compared to the model (blue). (H) Mean time of peak ICI in the distribution. (I) Example conversation showing paired and trio conversations. (J) The intercall interval in the data (pink) compared to base model. The number (K) and mean length (L) of the conversations. The (M) change in volubility and (N) model-corrected change in volubility of the high, medium and low caller in each three caller session session. (O) Difference in the proportion of all conversations that included only 2 out of the three monkeys. * p<0.05, ** p<0.01, *** p<0.001, ****p<0.0001

Overall, our model accurately captures the conversation dynamics of marmoset dyads, and that this is in a large part modulated by responses to individual calls. It is, however, important to consider that although the other factors did not significantly affect the ICI or conversations in this social context, they do reflect true dynamics of the behavior (e.g. a lack of interruptions) and may be more integral in more complex social contexts.

### Vocal behavior across different social contexts

The vocal turn-taking model shown above effectively captures the dynamics of marmoset dyads in a conversation. However, this is not the only context in which marmosets communicate, nor is it the only context in which marmosets engage in conversational turn-taking. While in some instances single marmosets may be isolated from the group, more commonly multiple marmosets are present in a scene during which vocal behavior represents a dynamic communication network. To test whether vocal behavior generalizes to other behavioral contexts, and how well our turn-taking model performs in these conditions, we tested marmosets in a separate set of experiments designed to characterize vocal behavior in single animals and trios of visually occluded individuals.

First, we determined whether the vocal behavior changed across the three paradigms (solo, paired and trios), by comparing 12 monkeys that were included in all paradigms. We saw no difference in the mean intercall interval between subsequent calls by a monkey (ICI self, Figure 4A), the response delay (Figure 4B), or call length (Figure 4C), although trends towards shorter and faster calls when more monkeys were interacting were present across the data. The monkeys did not change their calling rate across contexts (Figure 4D), and the calling rate in both the solo (Figure 4E) and trio (Figure 4F) conditions were highly correlated to the paired calling rate for individual monkeys, suggesting that individual differences in volubility were stable across these social contexts.

While no turn-taking behavior occurred in the solo condition, it is possible to quantify and model other aspects of their vocal behavior. We recorded a total of 77 solo session from 27 monkeys. Seventeen of these marmosets were also included in the two-monkey experiments described above. We used the model based on the paired monkeys to generate the calling dynamics in the solo sessions, resulting in an ICI distribution that was not significantly different (p=0.29, linear mixed-effects model, Figure 4G), suggesting our model also broadly captured the vocal behavior of single marmosets. However, a (non-significant) delay in peak ICI in the behavior data compared to the model was evident (Figure 4H, p=0.42). We hypothesize, based on the trends in the earlier data, that this may reflect an interruption avoidance strategy of the marmosets in the two monkey data^11,34^. Specifically, marmosets reduce their response delay and call length when other marmosets are present in the scene, which is reflected in the shorter peak ICI in the modelled data, based on the two-monkey recordings.

To further test whether our model generalized to more complex social scenes, we examined marmoset vocal behavior in a social context comprising multiple callers. Here we recorded the vocal behavior of three marmosets who were visually isolated from each other (See Methods). We recorded the vocal behavior of 21 marmosets across 25 sessions, 19 of these subjects were also included in the two-monkey experiment included above and used the model based on the paired monkeys described above to generate the calling dynamics in the three monkey sessions. The ICI distribution in multi-monkey context that was not significantly different between the two monkey contexts (p=0.15, linear mixed-effects model, Figure 4J). Neither the number of conversations (Figure 4K), nor the length of the conversations (Figure 4L), was significantly different from the model conversations.

Overall these results insinuate that marmosets do not change their vocal behavior depending on the social context, despite vocal interactions being modulated by social context across species^14,35–37^. It is, however, important to note that the model is based on the measured volubility during each session, so a session-wide effect on the volubility, whereby certain monkeys are suppressed while others increase their volubility may be present and not captured by these initial analyses. Indeed, when we measured the relative change in volubility of monkeys compared to their previously measured volubility in the solo and paired experiments, we show that on average one caller in each session significantly down-modulated their volubility (Figure 4M). This effect remained in the model-corrected volubility (see Methods), which corrects for the effects of the other monkeys’ measured volubility (Figure 4N). This suggests that a social suppression factor should be included in the model to account for scenes comprising multiple marmosets. We modelled a suppression factor for the non-conversation partner during ongoing conversations by the other individuals. We tested 9 levels of suppression (0-20% with 2.5% steps). Comparisons of these suppression models to the data showed that although no model captures all aspects, the 10% model shows the most similar proportion of dyads to the data, and shows the closest fit with the data overall (Figure 4O, SFig 5A-D). This suppression model exhibited an improvement over the base model, as the conversations it produces are more similar in structure to the data, suggesting a conversation-specific socially mediated reduction in volubility is present in multicaller groups. What remains unclear, however, is what determines which individuals suppress their calling at the outset of a session (i.e. low caller) and why the conversationalists change when they do. This highlights the fact that key sources of variance in this behavior have yet to be incorporated into our model and that targeted experimental work is likely needed to fully account for certain facets of marmoset vocal turn-taking.

**Figure 5:**
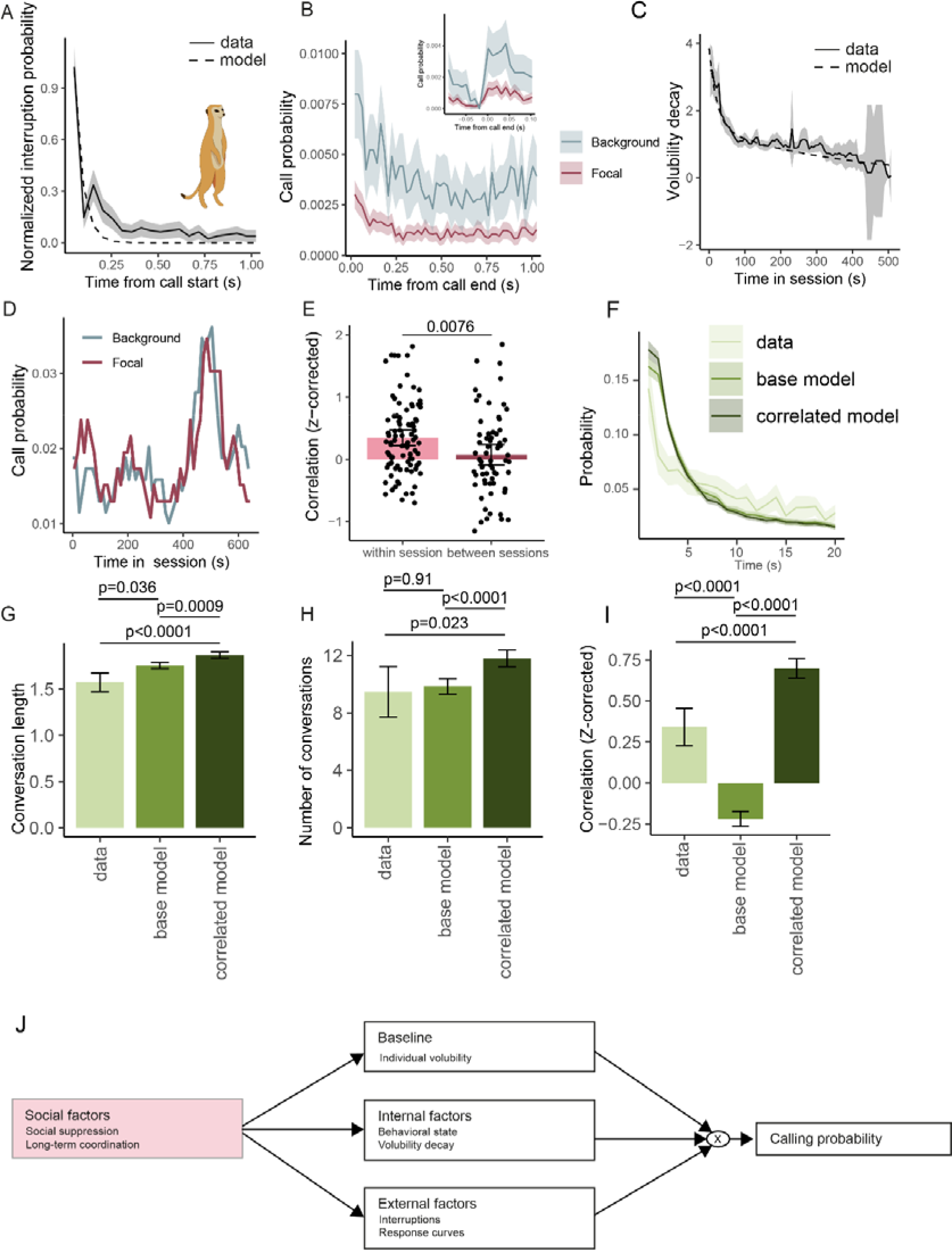
Vocal turn-takin model applied to meerkat vocal interactions. (A) Interruption curve, modelled by an exponential decay (dashed). (B) Call probability following a call by the partner, inset shows zoom of call probability around call end. (C) Volubility decay across sessions, modelled by an exponential decay (dashed). (D) Example of within session variability of focal and background meerkats. (E) Correlation of focal and background meerkats within and across sessions. (F) ICI, (G) conversation length, (H) number of conversations, and (I) correlation between the focal and background callers in the data, base model, and correlated model. (J) Schematic of novel model that includes a social effect on the base factors of the model.

## Vocal behavior modulation in social contexts across species

Next, we investigated whether the general rules that govern vocal interations in marmosets apply more broadly to other species. To this end, we analyzed a publically available dataset of multi-caller turn-taking in meerkats (*Suricata suricatta*) during sunning by Demartsev et al.^33,38^. In this elegant study, each session included a focal caller (one individual), and an unknown number of background callers. Although meerkat vocal interactions occur at a more rapid periodicity than marmosets, they are suggested to involve similar principles, such as interruption avoidance and response modulation^33^. We fitted the individual model components developed in the multi-caller essions from marmosets to these data.^33^ Similarly to marmosets, the interruption curve in meerkats follow an exponential decay (Figure 5A). Although the meerkats show an increased calling probability after a conspecific call (Figure 5B), the response curve did not possess an initial dip followed by peak as seen in the marmosets. Instead, the meerkats initiated their response immediately after the previous call ended (Figure 5B inset). The data included larger groups of meerkats, with few consecutive calls by the focal callers, resulting the self response curve to be dominated by calls that represent rapid back-and-forths between the focal caller and background callers (SFig 5E). Indeed, when only self responses without interjectors are included, no clear pattern arises in the self response curve, aside from a lack of calls in the first 0.25 s, which reflects the time delay between notes to be considered part of a single call^33^. The internal factors also follow the same principles as the marmosets, with an overall decay in calling over time (Figure 5C), as well as variability within sessions (Figure 5D). However, contrary to the marmosets (Figure 5S Fig 3B), the meerkats largely coordinate their session-wide variability (Figure 5E), suggesting their behavioral state is further modulated by a social factor that causes the animals within a group to increase their volubility at the same time.

We trained our model using the factors as described above, with the exception of the self-response, which we excluded apart from a suppression in the first 0.25s following a call, due to the low number of data points and high variability. As the identity of the background meerkats was not provided in the dataset, we treated them as a single conversation partner. We altered the time resolution of the model to account for shorter reponse latencies of meerkat interactions, and allowed for responses to start in the same time bin as the previous call end. Moreover, to determine whether the meerkats correlate their behavioral state, we tested an additional model where the initial behavioral state of both meerkats was identifical (‘correlated model’). We found an altered ICI in our model (Figure 5F), paired with an increased conversation length (Figure 5G), though or model did produce a comparable number of conversations (Figure 5H). The changes may be due to our model not allowing interruptions during a call, as well as the assumption of only 2 callers, while in reality multiple background callers will be present. Despite these differences we could show that the correlated behavioral state was not simply emergent from the caller dynamics, as shown by the lack of correlation in the base model (Figure 5I), suggesting the correlated behavioral state may represent coordination of the meerkats in this social context.

Together the tested multi-animal contexts show that while our model captures core processes for this vocal behavior across taxa, specific contexts cause additional socially-mediated effects that need to be included to fully describe the behavior across species and contexts (Figure 5J). Specifically, the marmoset vocal turn-taking model based on two monkey data more effectively captures marmoset behavioral dynamics in multi-caller environments if a suppression factor is included, while the meerkat data includes behavioral state modulation between animals. Importantly, our model demonstrates that while social factors significantly affect the dynamics of vocal turn-taking in meerkats, particularly in scenes comprising multiple conspecifics, the same core principles, including the interruption decay curve, response curves, and volubility decay, are consistent across species.

## Discussion

Here, we propose a new model of marmoset vocal turn taking. Leveraging the extensive analyses of marmoset vocal behavior in multiple behavioral contexts described here, we show that marmoset vocal turn-taking during conversations involves a suite of complementary processes whose collective – and respective – influence on this behavior had not collectively been considered previously. Although previous studies found varying effects of social and behavioral factors^21,28,39,40^, these have to date not been combined in a comprehensive computational model of marmoset vocal interactions. Our model balances individual differences in volubility with social and behavioral factors; the foundation of which is evidence showing that marmoset calling probability is modulated by several internal and external factors, including a volubility decay over time and the behavioral state, as well as the occurrence of a conspecific producing a call. More complex multi-speaker environments revealed that other social factors may affect these vocal interactions. Conversations, for example, bias to dyads and the conspecific in the scene who is not in a conversation will suppress their calling during the interaction. We furthermore demonstrate that another (non-primate) species follows the same principles, and our model can readily be adapted to investige vocal interactions in these species to identify shared and idiosyncratic processes that underlie this dynamic behavior across taxa. Our turn-taking model illustrates that multiple distinct factors underlie the dynamics vocal turn-taking across species and social contexts and provides a new framework for explicating its underlying mechanisms.

The model of marmoset turn-taking developed here revealed that at least four separable factors have differential effects on vocal behavior, providing a key foundation upon which to further elucidate this dynamic behavior in marmosets and across other species. Marmoset calling probability during conversations was affected by (1) an individual’s inherent volubility, (2) internal factors – i.e. state and volubility decay, (3) external factors – i.e. the behavior of the partner, and (4) social factors such as the occurrence of other conversations in the social scene. Importantly, the independent influence of these factors was not solely evident from behavioral analysis, but only became apparent when modelling each source of potential variance affecting marmoset conversational dynamics in different contexts. Because of the identification of these distinct factors, this model has implications to better understanding turn-taking more broadly. Particularly in species whose turn-taking behavior differs from marmosets. Meerkats^33^, as shown, engage in conversational turn-taking of three or more individuals, unlike the bias to dyads in marmosets, while singing mice^32^, naked mole-rat^41^, and bushcricket^42^ turn-taking occurs at significantly faster intervals than is observed in marmosets. Other species, such as elephants^36^, alter their interruption rate based on the conversation partner. As demonstrated here by applying our model to meerkats, this framework can be explicitly used to identify shared processes underlying turn-taking across taxa as well as unique features of the behavior. Accounting for these similarities and differences across species in our model would potentially make key predictions about the underlying mechanisms and by extension a path for the elucidating the evolution of the behavior.

One notable limitation of our study is that we could not account for all sources of variance in the vocal behavior. While some elements of marmoset conversations appear stochastic, they may simply not yet be accounted for by the model. The addition of further behavioral quantification, such as body posture, overall arousal, and more detailed analysis of state fluctuations can further refine the model and increase its explanatory power. As occurred following analysis of the multi-caller contexts, the model can be updated with additional factors to account for such discrepancies and describe the underlying dynamics in more detail. In addition, as our model is stochastic in nature, it has limited predictive power for the timing of individual calls in conversations, but rather simulates new conversations that match the data in overall structure. Although the probability calculated by the model may be used to predict responses over short time periods, it does not predict session-wide conversations. Lastly, although our model accurately describes the contexts tested here, it does not yet account for both the rapid^15,34^ and structural^11^ adaptations the monkeys are able to make in the presence of interfering noise. This behavior is likely crucial to their vocal communication, as it is needed to effectively communicate in an environment with competing background noise^43^.

The factors included in our vocal-turn taking model are based on analysis of the behavioral phenotype, but they represent the neural mechanisms, or suites of mechanisms, that support marmoset conversations. The model is based on a probability measure, calculated using the base probability and the various aforementioned factors. This probability-based approximation may manifest as neural activity with random variability crossing a certain threshold to initiate a call, reminiscent of decision-making dynamics.^20,44,45^ Several prefrontal cortex (PFC) regions have been indicated in decision making studies^46^, as well as anterior cingulate cortex (ACC)^46–48^. Given its role in vocal control^49^, this area is a promising candidate for further studies into the neural correlates of marmoset volubility and suppression of calling in multi-animal contexts. The model furthermore includes a response curve, which represents an increased probability of calling following a vocalization by the partner. We hypothesize such a vocalization may represent a cue signal affecting ventrolateral PFC,^49,50^ leading to a ramping of activity, either in this area, or downstream in ACC, which ultimately causes the monkey to produce a response. Indeed, recent studies have shown that such ramping following a cue signal indeed occurs in the anterior cingulate cortex in macaques,^49^ although other prefrontal areas may additionally be involved.^51^ The turn-taking model described here provides a powerful framework to better understand the processes affecting this natural vocal behavior and a roadmap for explicating the neural mechanisms that underlie each of these factors, both in these marmosets and more broadly for comparative analysis of other species.

## Acknowledgements

This work was supported by NIH grant R01 R01 DC012087 to CTM. All research was approved by the UCSD Institutional Animal Care and Use Committee.

## Author Contributions

DMG and CTM conceived of the study and wrote the manuscript. DMG and DAF collected the data. DMG performed the formal analyses, data curation and visualization of the results. CTM supervised the study and acquired the funding.

## STAR Methods

### Resource availability

3 Resource availability
| REAGENT or RESOURCE | SOURCE | IDENTIFIER |
| --- | --- | --- |
| Deposited data |  |  |
| Data and Software | Dryad | https://doi.org/10.5061/dryad.9ghx3ffpx |

### Lead contact

Further information and requests for resources should be directed to and will be fulfilled by the lead contact, Cory Miller

### Materials availability

This study did not generate any new materials or reagents.

### Data and code availability

The preprocessed data supporting this study are available on the Dryad public repository. The raw audio files are not available in a public repository because of the size of the files but are available from the lead contact upon request.

### Experimental model and subject details

*Subjects*. A total of 53 unique subadult and adult marmosets were used across the three experiment. For the two monkey experiments, a total of 42 unique monkeys were included (48% female), with ages between 284 and 2737 at the time of recording. For the single monkey experiments 27 unique monkeys were included, of which 17 were also included in the two monkey recordings (59% female), with ages between 331 and 3421 days at time of recording. For the three monkey experiments 21 unique monkeys were included, of which 19 were also included in the two monkey recordings (48% female), with ages between 572 and 2466 days at time of recording. All animals were group housed, and experiments were performed in the Cortical Systems and Behavior Laboratory at University of California San Diego (UCSD). All experiments were approved by the UCSD Institutional Animal Care and Use Committee.

## Method Details

### Experimental setup

All recording sessions took place in a Radio-Frequency shielding room (ETS-Lindgren) in a 4x3x3 m room. Animals were placed in a 32x18x46cm box, with a mesh on one side, and placed on a table on either side of the room. For the three monkey experiments, a third table was added such that the three table formed a triangle. All boxes were separated with an opaque black curtain. Each box had a directional microphone pointed towards it (Sennheiser model MKE 600 and ME-66), which was amplied using a preamplifier (PreSonus BlueTube DP v2). Data from all microphones was acquired using a custom MATLAB script.

## Quantification and Statistical analysis

### Coupled oscillator analyses

First we calculated the interruption rate by determining the number of overlapping calls. We obtained the shuffled dataset by randomly combining 2 monkeys from different sessions and calculating the interruption rate for these shuffles. The phase response curve (PRC) was calculated as previously described^8^, in short, we calculated the response interval (R, mean time between a call and a response), phase response (PR, mean time between two calls of the same monkey, when partner called in between) and median call interval (T0, mean time between two calls of the same monkey with no calls in between) for each session. We included the bias correction points, which are any calls where a monkey made two consecutive calls, and the partner responded after the second call (i.e. PR<R). We calculated the mean T0 across all sessions and then determined the PRC correlation both before and after T0. Next, we randomly selected subsets of sessions of various sizes and performed the same PRC analyses on these subsets, to determine the effect of the number of datasets included on the PRC correlation. We calculated the PRC and p-value using the corrcoef function in MATLAB.

### Predictive models

We first extracted 12 predictive factors for each call, most representing characteristics of the preceding call, and inspired by previous work^22^:

‐ Monkey calling rate: The mean cumulative (i.e. only including calls made before the call in question) calling rate of the monkey producing the call.
‐ Call # in conversation: How many consecutive (within 30 s) calls have been made so far in the current conversation
‐ Maximum frequency: The maximum frequency of the call
‐ Loudness: The relative loudness of the call, normalized to the mean loudness of all calls produced by the same monkey in that session
‐ Number of pulses: The number of phee pulses the call consisted of
‐ Ending slope: The slope from peak frequency to phee offset (Hz/s) of the call.
‐ Start frequency: The frequency at phee onset of the call
‐ Length of the last pulse: The length (in s) of the last phee pulse of the call.
‐ Partner call length: The total length (in s) of the call.
‐ Partner calling rate: The mean cumulative (i.e. only including calls made before the call in question) calling rate of the partner monkey.
‐ End frequency: The frequency at phee offset of the call.
‐ Partner ICI: The response delay of the call (i.e. between the last call of the partner monkey and the current call).

We fitted generalized linear mixed-effects models using the *glmer* function in R, with the 12 factors as the fixed-effects, and the monkey and partner ID as the random effects, to obtain the odds ratio. Prior to fitting the linear model and random forest, we counterbalanced the data by including an equal number of responses and non-responses per session. We fitted a linear model (function *fitclinear* with crossvalidation on) on this data to predict whether the partner would respond to this call, and calculated the prediction error to determine the accuracy. The random forest was fit using the *TreeBagger* function in MATLAB, with the predictor selection set to *‘interaction-curvature’*, and accuracy was again obtained by calculating the prediction error. We extracted the predictor importance in the linear model from the absolute weights of the predictors, and from the random forest using the built-in *OOBPredictorImportance*.

### Marmoset phee communication model

#### Internal state model

First we calculated the rolling average calling rate in 55 180 s windows with a step size of 30 s. We used 5 consecutive windows as the predictor with the subsequent window as the dependent variable in our model. We trained a linear model using the built-in *fitlm* function in MATLAB. We extracted the prediction error of the model and fit a Gaussian distribution using the fitdist function. Next, to model new internal state values, we randomly selected 5 values from a Gaussian distribution fit on the calculated mean calling rates. We then recursively predicted the subsequent value using the linear model, and added a random error to this value drawn from the Gaussian distribution fit on the prediction error. We threw out the first 70 values, to prevent any effect from the initial 5 seeding values, and divided the remaining internal state values into 200 sessions of 70 values, each normalized to have a mean of 1. For each modelled monkey, an internal state vector was randomly drawn, and the vector was interpolated (*interp1* function, using *pchip* interpolation) to obtain an internal state value per second. To prevent edge effects from the interpolation and correlations caused by the continuous nature of the initial generation of the internal state, the interpolated values consistent of the values outside of the 1800s session length were dropped.

#### Arousal

We used the same rolling average calculated for the internal state to obtain the mean calling rate across each session. These were averaged across all sessions and monkeys to obtain a single vector, which was normalized to have a mean of 1. We fitted this average with a two-term exponential decay model interpolated (*fit* function, using *exp2* fittype), and used the resulting model to generate the arousal decay over time for each session.

#### Interruptions

The interruption curve was calculated by determining the mean number of calls by the partner monkey while a monkey was vocalizing, calculated in 0.2s windows for a total of 5 s. This mean number was normalized to 0-1 range, and fit with an exponential decay with an origin of 1, using the MATLAB fit function.

#### Response curve

We calculated the mean response curve by determining the mean number of calls by the caller and by the partner occurring in the 30s following the end of a call. This response curve was then smoothed for each monkey and session using the *smooth* functon with a local regression using weighted linear least squares and a 2nd degree polynomial model. The resulting smoothed response curves were then normalized to the base rate, defined as the response likelihood in the last 10 seconds of the 30s curve, and averaged to obtain the response profile.

#### Correction factor

As the model varied the probability of calling depending on the internal state and occurrence of calls, a discrepancy in the inputted calling rate and the resulting modelled rate occurred. The size of this discrepancy depended on the inputted calling rates, and as such, we modelled a correction factor to account for this. We first built the model as specified, and then generated model conversations for input calling rates of 0-300 calls in 10 call steps for each monkey, and calculated the discrepancy. We then fit a linear model (*fitlm* function) to predict the discrepancy in calling rate based on the monkey and the partner calling rate. We used this correction model to correct the calling rates prior to running the model, so it generated the same number of calls as the inputted values for each monkey.

#### Model-corrected calling rate

Using the linear model for the correction factor, we can calculate the original input calling rate required to produce the output calling rates observed. As the correction is differentially affected by both the caller and its partner’s rate, we calculate the correction for both separately (using the same linear model) to obtain the model-corrected calling rate.

#### Conversation modelling

Each 1800 s session was divided into 0.2 s bins, and we determined a baseline probability for each bin by multiplying the calling rate of each monkey, the correction factor calculated for each monkey, the arousal curve and the internal state curves randomly selected for each monkey. Next, we iterated through each timepoint (i.e. 1800 x 5 timepoint total), and drew a random number for each monkey (MATLAB *rand* function). If this number was smaller than the probability of calling at that timepoint, a call occurred for that monkey. The call length was randomly drawn from a Gaussian distribution fitted to the actual call length. The probability for the subsequent time bins was altered by multiplying the appropriate interruption curves and response

#### curves with the existing probability

We performed this action iteratively until the end of the session was reached, and outputted the list of calls and who made each call for further analysis.

#### Cross-validation

As the model was based on the 2 monkey data, we applied cross-validation to generate the 2 monkey data sets. The total number of datasets (N=107) was divided into 5 folds. We then fitted the four factors on all but one fold, and used the resulting factors and curves to generate the model data for the 5^th^ fold. As such, the modelled session was never included in the model data.

### Meerkat communication model

We obtained the freely-available meerkat calling data from Demartsev et al.^33,38^ to build the meerkat communication model. The meerkat model used the same procedure as described above with some notable modifications to better fit the timescale and nature of the data:

‐ The timesteps were reduced to 0.05s (compared to 0.5s in the marmosets).
‐ The self response curve prohibited self-responses in the first 0.25s following a call, to fit with the definition of individual calls, but did not modulate the calling further.
‐ Individual responses could take place in the same time bin as the end of a call, to allow for the short timescale of the meerkat calls.

## Supplementary figures

**S Fig 1.**
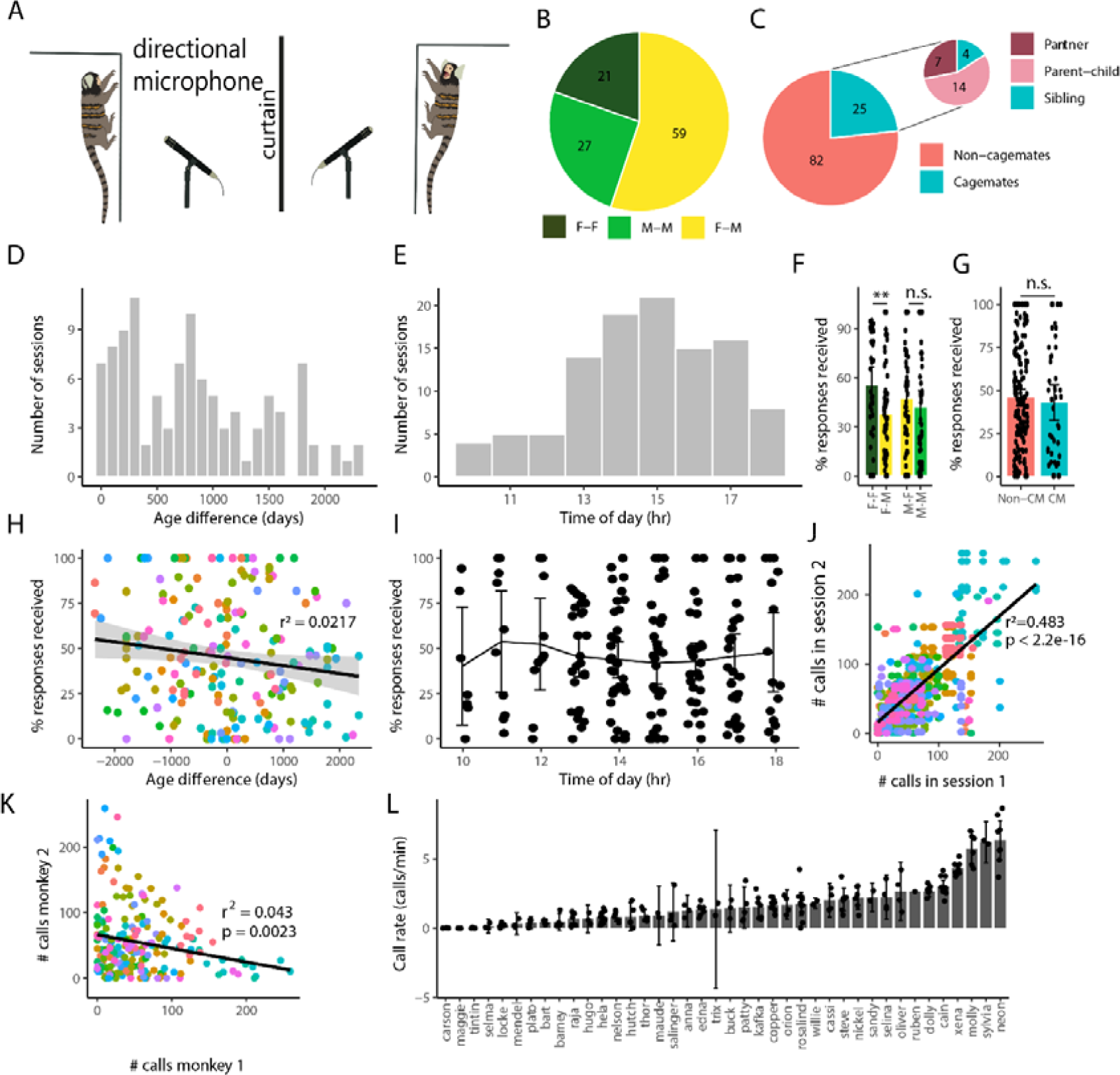
Marmoset response rates across environmental and social contexts. (A) Schematic of the experimental setup, with two marmosets separated by an opaque curtain, and a directional microphone in front of each monkey. (B) Distribution of female-female (dark green), male-male (light green) and female-male (yellow) sessions. (C) Distribution of the cage mate relations included in our dataset. (D) Absolute difference in age of the two monkeys for each session. (E) Time of day the session was recorded. (F) Percentage responses received as a function of the sex of the monkeys. (G) Percentage of calls responded to as a function of the cage mate (CM) status. (G) Percentage responses received as a function of the age differences between the two monkeys included in the session (H) Percentage of calls receiving a response as a function of the time of day. (J) Correlation between the calling rate of between sessions of the same monkey. (K) Correlation of calling rate between the two monkeys per session. (L) Calling rate of each monkey included in the dataset. Errorbars show 95% CI.

**S Fig 2:**
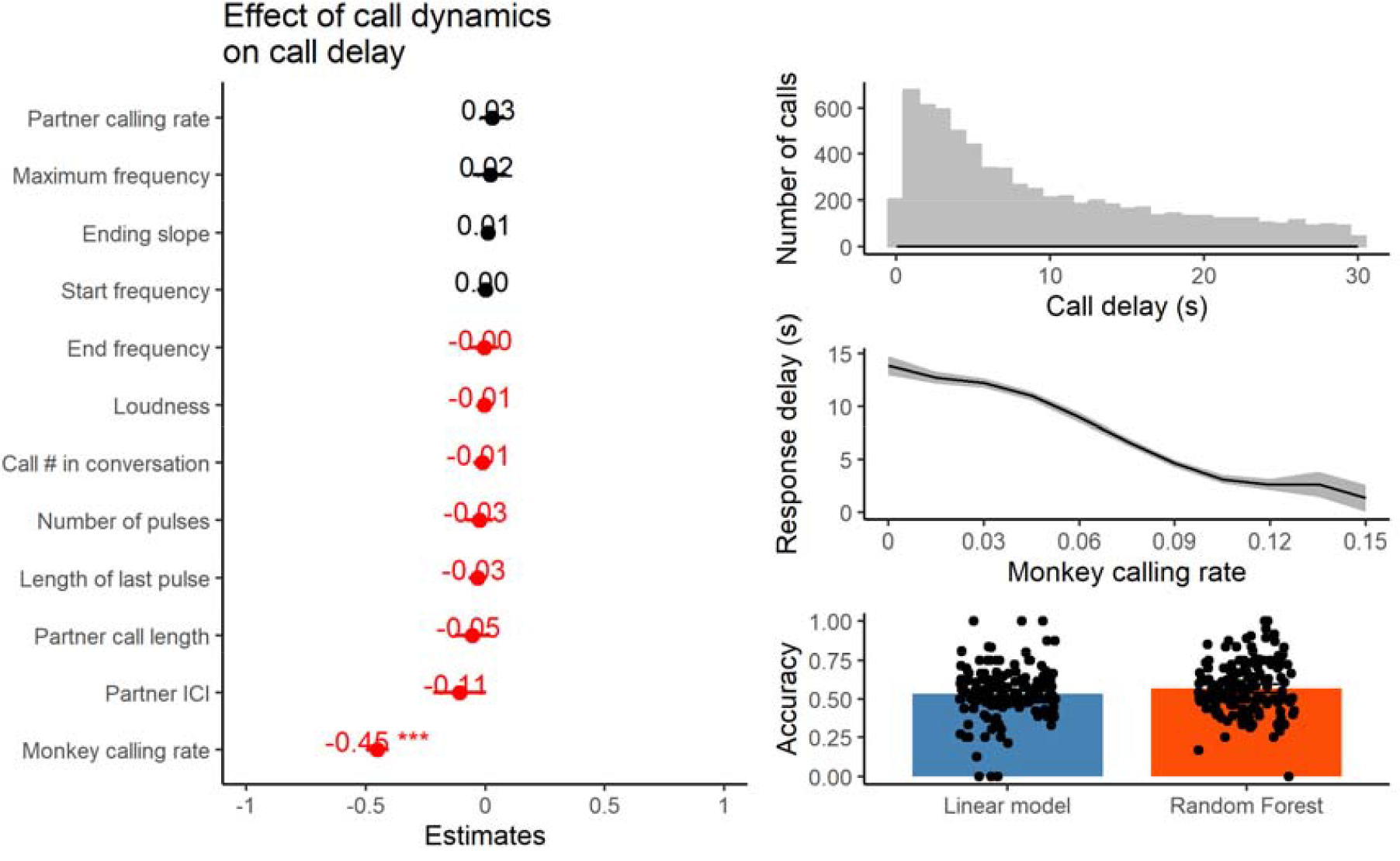
Effect of various call parameters on call delay. (A) the estimated effect size of the parameters, as calculated from the Linear Mixed-effects model. (B) The overall distrubtion of call delay. (C) The call delay as a function of the monkey’s calling rate. (D) The accuracy in prediction the response likelihood of a linear model and a random forest, when monkey calling rate was excluded.

**S Fig 3.**
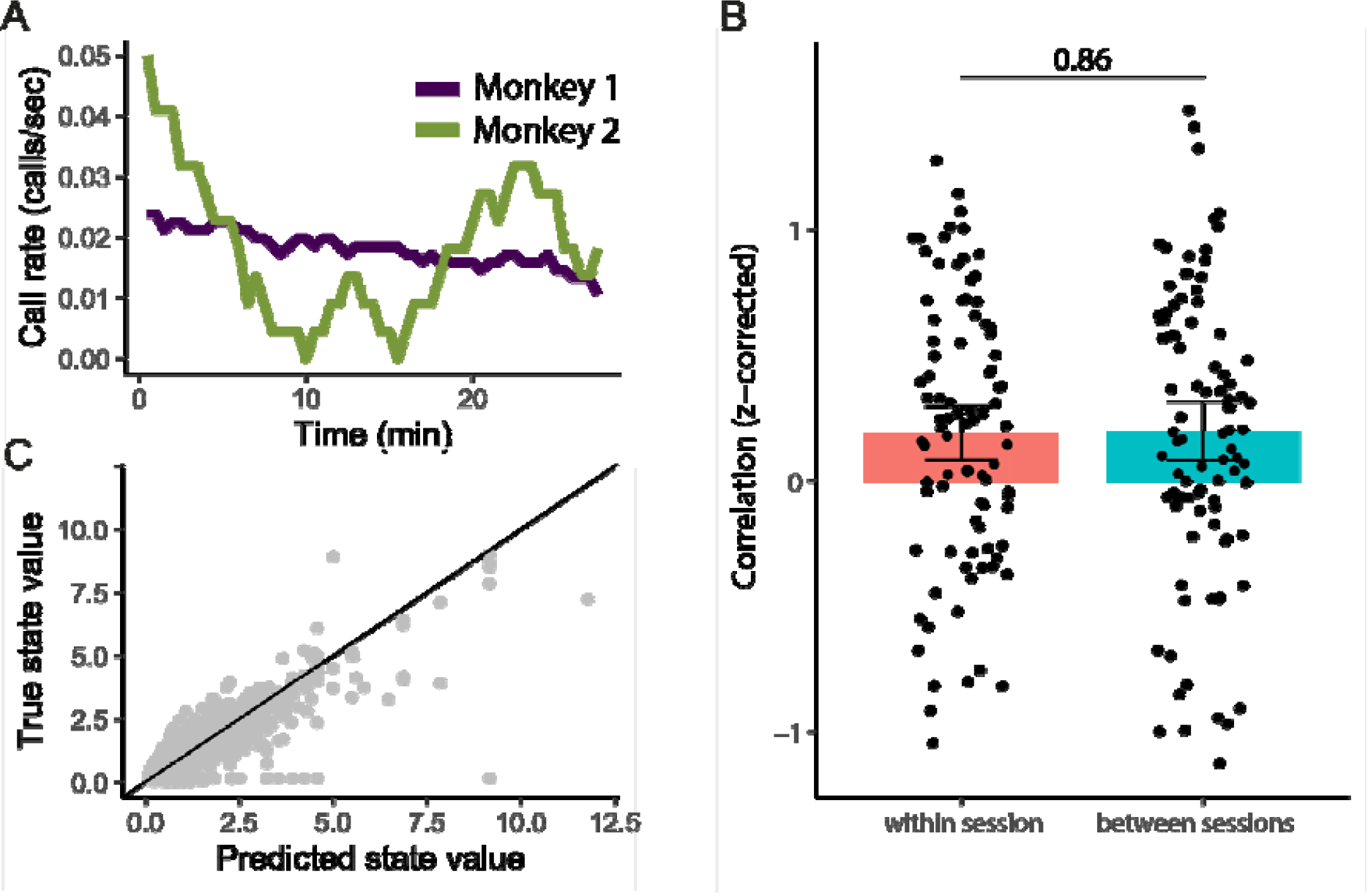
State modulation of marmoset calling rate. (A) Example calling rate across a single session of a pair of monkeys. (B) Correlation of the calling rate between the pairs of monkeys (pink), or a shuffle control of pairs of monkeys from different sessions (blue). (C) Performance of the linear model showing the predicted state value (x-axis), compared to the true state value (y-axis), the black line indicates a perfect score.

**S Fig 4.**
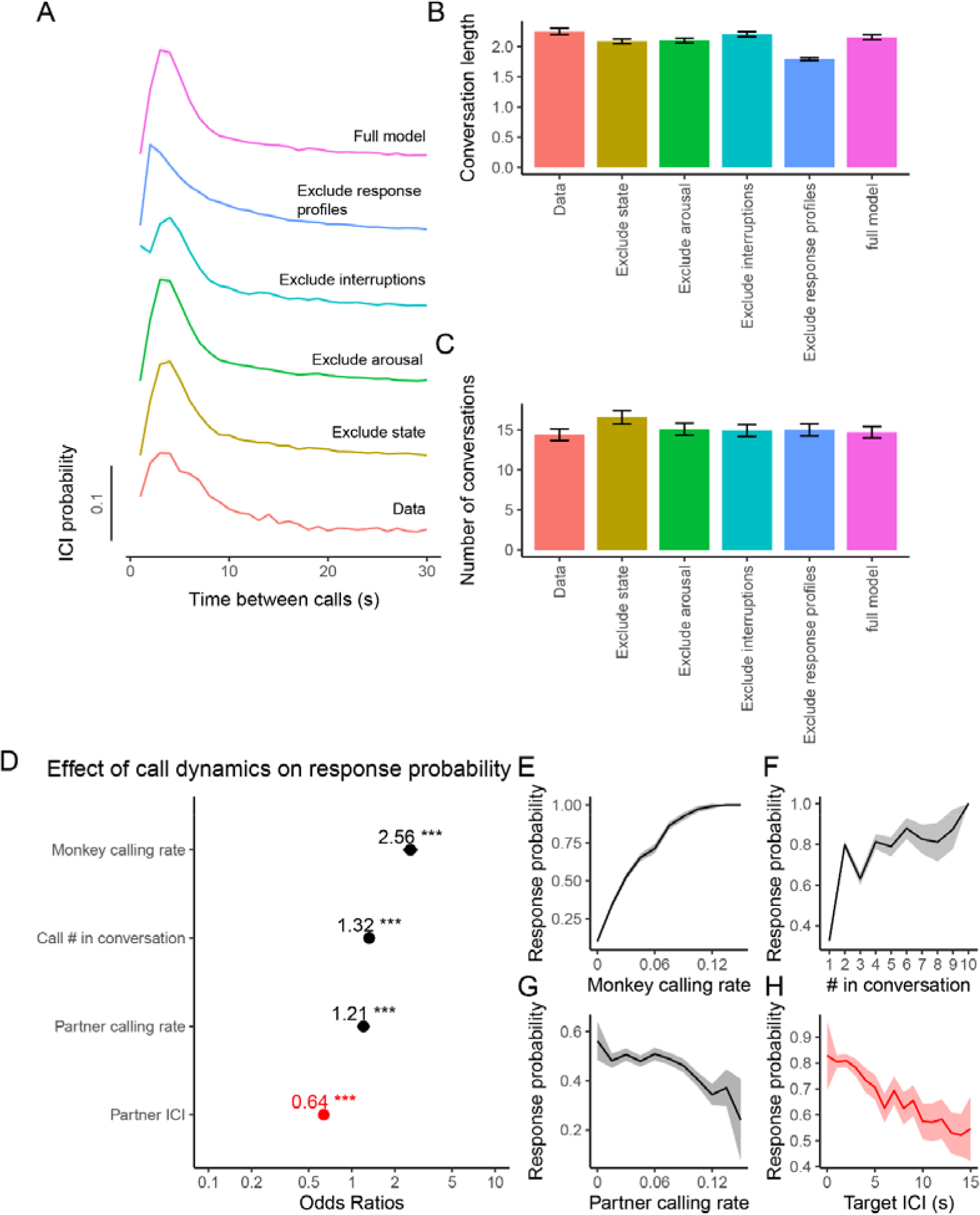
Impact of each individual factor on ICI and conversations. (A) The ICI curve when each of the 4 different factors in the model was excluded. The conversation length (B) and number of conversation (C) when each of the factors was excluded. (D) Odds ratio of 4 parameters as resulting from the Generalized Linear Mixed-effects Model on modelled data. Effect of (E) monkey calling rate, (F) position of call in the conversation, (G) partner calling rate, and (H) the target ICI on the response probability

**S Fig 5.**
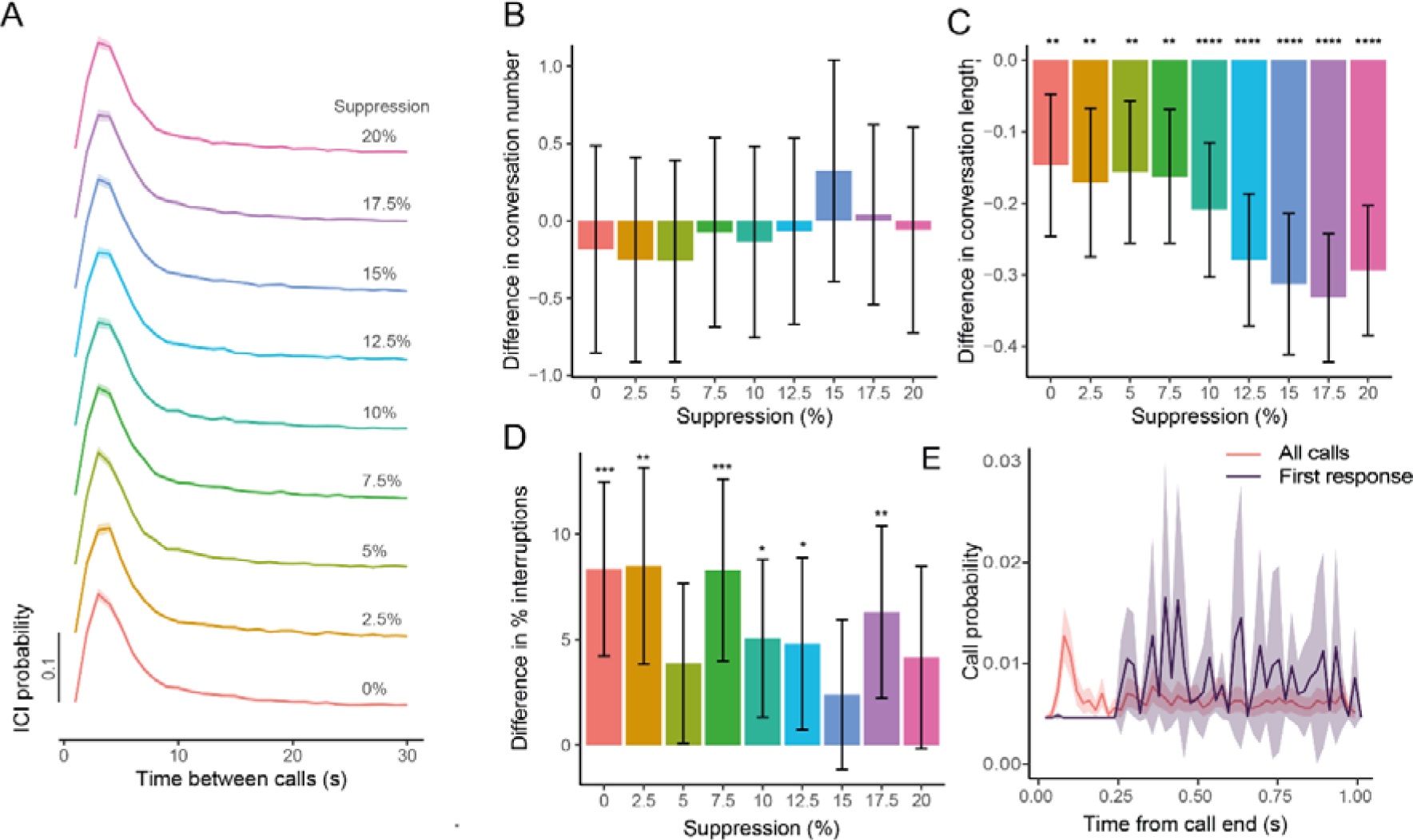
Calling dynamics for three monkey experiments. (A) ICI curves, (B) difference in conversation number, (C) difference in conversation length, and (D) difference in % interruptions by the non-conversation partner compared to the data for the models with varying levels of suppression of the non-conversation partner. (E) Self-response curve when using all calls (pink), or when only including calls where the first response is by the focal caller (purple). * p<0.05, ** p<0.01, *** p<0.001, ****p<0.0001

